# Advanced lesion symptom mapping analyses and implementation as *BCBtoolkit*

**DOI:** 10.1101/133314

**Authors:** C Foulon, L Cerliani, S Kinkingnéhun, R Levy, C Rosso, M Urbanski, E Volle, M Thiebaut de Schotten

## Abstract

**Background:** Patients with brain lesions provide a unique opportunity to understand the functioning of the human mind. However, even when focal, brain lesions have local and remote effects that impact functionally and structurally connected circuits. Similarly, function emerges from the interaction between brain areas rather than their sole activity. For instance, category fluency requires the association between executive, semantic and language production functions.

**Findings:** Here we provide, for the first time, a set of complementary solutions to measure the impact of a given lesion upon the neuronal circuits. Our methods, which were applied to 37 patients with a focal frontal brain lesion, revealed a large set of directly and indirectly disconnected brain regions that had significantly impacted category fluency performance. The directly disconnected regions corresponded to areas that are classically considered as functionally engaged in verbal fluency and categorization tasks. These regions were also organized into larger directly and indirectly disconnected functional networks, including the left ventral fronto-parietal network, whose cortical thickness correlated with performance on category fluency.

**Conclusions:** The combination of structural and functional connectivity together with cortical thickness estimates reveals the remote effects of brain lesions, provide for the identification of the affected networks and strengthen our understanding of their relationship with cognitive and behavioural measures. The methods presented are available and freely accessible in the *BCBtoolkit* as supplementary software [1].

---

Recent advances in neuroimaging techniques, allowed for the further examination of the structural and the functional organization of the human brain. While diffusion weighted imaging (DWI) tractography [2] depicts how brain areas are connected together, functional magnetic resonance imaging (*f*MRI) measures the activity within and interaction between brain areas in the elaboration of functions [3]. These methods have been successfully applied to the healthy human brain, however, they remain underused in patients with brain lesions.

Patients with brain lesions provide a unique opportunity to understand the functioning of the human mind. Lesion symptom mapping analyses traditionally assume that visible and directly damaged areas are responsible for a patient’s symptoms [4-7]. Following this logic, the areas that are the most frequently damaged by the lesion are considered as the neuronal substrate for the function. Previous studies employing this method have pinpointed critical areas dedicated to, for example, language production [8], comprehension [9], spatial awareness [10-13] and other high-level cognitive functions [14-17]. However, anatomical disconnections between regions are also important considerations for the exploration of cognitive deficit [18, 19]. The dysfunction of distant areas that are connected to the lesioned tissue has also been reported in *f*MRI studies. They have shown that the networks are disrupted even by distant lesions through disconnection and diaschisis mechanisms [20-22].

Non-local effects of lesions have previously been explored using various forms of atlas-based analyses of tract damage [23-32], lesion-driven tractography [32-34], disconnectome-mapping [35-39] and lesion-driven resting state *f*MRI (rs-*f*MRI) connectivity [34, 40]. However, determining what these methods actually measure and identifying how to properly combine them are not always fully clear to the scientific community. Furthermore, there is an extremely limited availability of free, open-source software that applies methods to measure the non-local effects of lesions. These resources and scientific tools remain very much inaccessible and present a potential threat to reproducible science [41].

Disconnections and diaschisis can have an impact upon distant regions in several respects through maladaptive responses and pathological spread [42]. When disconnected from its inputs and outputs, a region can no longer contribute to the elaboration of the supported function. This phenomenon is called diaschisis [20, 21, 43]. Once deprived from its inputs and/or outputs, transneuronal degeneration in the region will occur [42], dendrites and synapses density will decrease in number, myelin content will be altered and neurons will reduce in size or die through a mechanism called apoptosis, a programmed cell death [44-46]. Hence, a white matter disconnection leads to both functional and anatomical changes that extend well beyond the visible damage. New approaches are therefore required to capture the long-range effects that follow brain disconnections. For instance, cortical thickness [e.g. 47] and other volumetric [e.g. voxel based morphometry 48] analyses have been previously used to study the structural changes associated with brain lesions, but have not been applied in the context of brain disconnection.

In response to this need, we provide here a set of complementary solutions to measure both the circuit, and the subsequent changes within the circuit that is caused by a lesion. We applied these methods to 37 patients with a focal brain lesion following a stroke or a surgical resection. We first assessed the risk of disconnection in well-known white matter tracts and tested their relationship with category fluency performance. Category fluency is an appropriate test to explore disconnection since it requires the association between executive, semantic and language production functions [49, 50]. We then developed a tractography-based approach in order to produce maps of the areas that are directly disconnected by the lesion and tested their relationship with category fluency performance. We additionally calculated the rs-*f*MRI connectivity of these areas to reveal the whole network of directly and indirectly connected regions that participate in category fluency. Finally, we explored potential microstructural changes in the latter disconnected regions, by estimating neuronal loss or local connectivity degeneration derived from MR-based measures of cortical thickness and resting state *f*MRI entropy.

## Methods

### Participants and Category fluency task

Thirty-seven right-handed patients (French-native speakers; 19 females; mean age 48 ±14.2 years, age ranging from 23 to 75 years) who presented with a frontal lobe lesion at the chronic stage (> 3 months) were included in this study (see table 1 for demographics). These patients were recruited from the stroke unit and the neuroradiology department at Salpêtrière Hospital, the neurological unit at Saint-Antoine Hospital and the neuroradiology department at Lariboisière Hospital in Paris. Patients with a history of psychiatric or neurological disease, drug abuse, or MRI contraindications were not included. Additionally, we gathered behavioural data from 54 healthy participants (French-native speakers; 27 females; mean age 45.8 ±14.4 years, age ranging from 22 to 71 years) in order to constitute a normative group. All participants performed a category fluency task [51] in French. They were instructed to enumerate as many animals as possible during a timed period of 120 seconds. The results were recorded by a clinical neuropsychologist (M.U.). Repetition and declination of the same animal were not taken into account in the final category fluency score.

**Table 1:** Demographical and clinical data

| ID | Age (years) | Education (years) | Gender | Lesion side | Lesion volume (mm<sup>3</sup>) | Lesion delay (months) | Aetiology |
| --- | --- | --- | --- | --- | --- | --- | --- |
| P01 | 56 | 17 | F | right | 255 | 7 | stroke |
| P02 | 55 | 19 | M | left | 34374 | 76 | hematoma |
| P03 | 46 | 17 | F | left | 14847 | 126 | stroke |
| P04 | 50 | 11 | F | left | 110145 | 137 | surgery |
| P05 | 64 | 14 | M | right | 59048 | 119 | stroke |
| P06 | 32 | 16 | F | right | 15946 | 129 | epilepsy |
| P07 | 51 | 11 | M | bilateral | 113170 | 54 | stroke |
| P08 | 70 | 5 | F | left | 51530 | 85 | surgery |
| P09 | 47 | 11 | M | right | 7809 | 115 | hematoma |
| P10 | 62 | 13 | F | bilateral | 21295 | 14 | hematoma |
| P11 | 41 | 16 | M | right | 55848 | 29 | surgery |
| P12 | 46 | 12 | M | bilateral | 2542 | 51 | hematoma |
| P13 | 67 | 15 | M | left | 4102 | 133 | stroke |
| P14 | 49 | 9 | M | bilateral | 14929 | 19 | hematoma |
| P15 | 36 | 14 | F | right | 40854 | 82 | surgery |
| P16 | 40 | 22 | F | left | 24829 | 56 | hematoma |
| P17 | 40 | 14 | M | bilateral | 14364 | 7 | hematoma |
| P18 | 23 | 16 | F | right | 21681 | 47 | surgery |
| P19 | 54 | 22 | M | right | 51897 | 48 | stroke |
| P20 | 71 | 17 | M | left | 25779 | 91 | hematoma |
| P21 | 23 | 15 | F | right | 29513 | 36 | surgery |
| P22 | 27 | 9 | F | left | 12986 | 30 | surgery |
| P23 | 26 | 13 | F | left | 2640 | 19 | surgery |
| P24 | 32 | 14 | F | left | 12653 | 4 | surgery |
| P25 | 59 | 16 | F | left | 97 | 9 | hematoma |
| P26 | 26 | 13 | F | left | 26928 | 32 | stroke |
| P27 | 58 | 12 | M | left | 1026 | 3 | stroke |
| P29 | 75 | 12 | F | left | 14938 | 16 | hematoma |
| P30 | 52 | 13 | F | right | 11978 | 20 | surgery |
| P31 | 58 | 12 | M | right | 13263 | 21 | surgery |
| P32 | 62 | 5 | M | right | 20281 | 9 | surgery |
| P33 | 41 | 17 | M | left | 7463 | 29 | surgery |
| P34 | 42 | 17 | M | left | 24319 | 6 | Infection |
| P35 | 60 | 12 | M | right | 41897 | 24 | surgery |
| P36 | 51 | 14 | F | right | 39213 | 17 | surgery |
| P37 | 51 | 12 | F | right | 8133 | 48 | surgery |
| P38 | 33 | 17 | M | right | 140947 | 48 | surgery |

The experiment was approved by the local ethics committee (Comités de protection des personnes, CPP Ile de France VI, Groupe hospitalier Pitie Salpetriere, Reference project number 16-10); all participants provided written informed consent in accordance to the Declaration of Helsinki. Participants also received a small indemnity for their participation.

### Magnetic resonance imaging

An axial three-dimensional magnetization prepared rapid gradient echo (MPRAGE) dataset covering the whole head was acquired for each participant (176 slices, voxel resolution = 1 × 1 × 1 mm, echo time = 3 msec, repetition time = 2300 msec, flip angle = 9°).

Additionally, the same participants underwent an *f*MRI session of resting state. During the resting state session, participants were instructed to relax, keep their eyes closed but to avoid falling asleep. Functional images were obtained using T2-weighted echo-planar imaging (EPI) with blood oxygenation level-dependent contrast using SENSE imaging an echo time of 26 msec and a repetition time of 3000 msec. Each dataset comprised 32 axial slices acquired continuously in ascending order covering the entire cerebrum with a voxel resolution of 2 x 2 x 3 mm. 200 volumes were acquired using these parameters for a total acquisition time of 10 minutes.

Finally, diffusion weighted imaging was also acquired for 54 participants of the normative group (French-native speakers; 27 females; mean age 45.8 ±14.4 years, age ranging from 22 to 71 years) and consisted in a total of 70 near-axial slices acquired using a fully optimised acquisition sequence for the tractography of diffusion-weighted imaging (DWI), which provided isotropic (2 × 2 × 2 mm) resolution and coverage of the whole head with a posterior-anterior phase of acquisition. The acquisition was peripherally-gated to the cardiac cycle [52] with an echo time = 85 msec. We used a repetition time equivalent to 24 RR (i.e. interval of time between two heart beat waves). At each slice location, 6 images were acquired with no diffusion gradient applied. Additionally, 60 diffusion-weighted images were acquired, in which gradient directions were uniformly distributed on the hemisphere with electrostatic repulsion. The diffusion weighting was equal to a b-value of 1500 sec mm^−2^.

### Stereotaxic space registration

As spatial normalisation can be affected by the presence of a brain lesion, additional processing was required before calculating the normalisation. For instance, in the case of bilateral lesions, the registration was weighted as previously reported [53]. For unilateral lesions, the first step was to produce an enantiomorphic filling of the damaged area [54]. Each patient’s lesion (or signal abnormalities due to the lesion) was manually segmented (using FSLview; http://fsl.fmrib.ox.ac.uk). Unilateral lesions were replaced symmetrically by the healthy tissue of the contralateral hemisphere. Enantiomorphic T1 images were fed into FAST [55] for estimation of the bias field and subsequent correction of radiofrequency field inhomogeneity. This improved the quality of the automated skull stripping performed using bet [56] and the registration to the MNI152 using affine and diffeomorphic deformations [57]. The original T1 images (non enantiomorphic) were registered to the MNI152 space using the same affine and diffeomorphic deformations as calculated above. Subsequently, lesions were segmented again in the MNI152 space under the supervision of an expert neurologist (E.V.). This method has been made freely available as the tool *normalisation* as part of *BCBtoolkit* [1].

The following sections of the manuscript are hypotheses-driven and outlined in supplementary figure 1.

### White matter tracts disconnection

Each patient’s lesion was compared with an atlas of white matter tracts [58], indicating for each voxel, the probability of finding a white matter tract such as the arcuate fasciculus, the frontal aslant tract or the uncinate fasciculus in the MNI152 coordinate system. We considered a tract to be involved when the likelihood of a tract being present in a given voxel was estimated above 50% [23]. This method is freely available as *tractotron* in *BCBtoolkit* [1]. We focused on frontal lobe tracts with a potential effect on executive, semantic and language functions since all of the patients had a frontal lesion. These tracts included the cingulum, the frontal aslant and the frontal superior and inferior longitudinal tracts for the executive functions [59], the uncinate and the inferior fronto-occipital fasciculi for the semantic access [60, 61] and the anterior and long segment of the arcuate fasciculi for the phonemic system [62, 63]. A Kruskall-Wallis test was employed to compare performance on the category fluency test for each tract between both preserved and disconnected patients and control participants. Subsequently, for each significant tract between patients, Mann-Whitney post-hoc comparisons were performed (**Fig.1**).

**Fig. 1:**
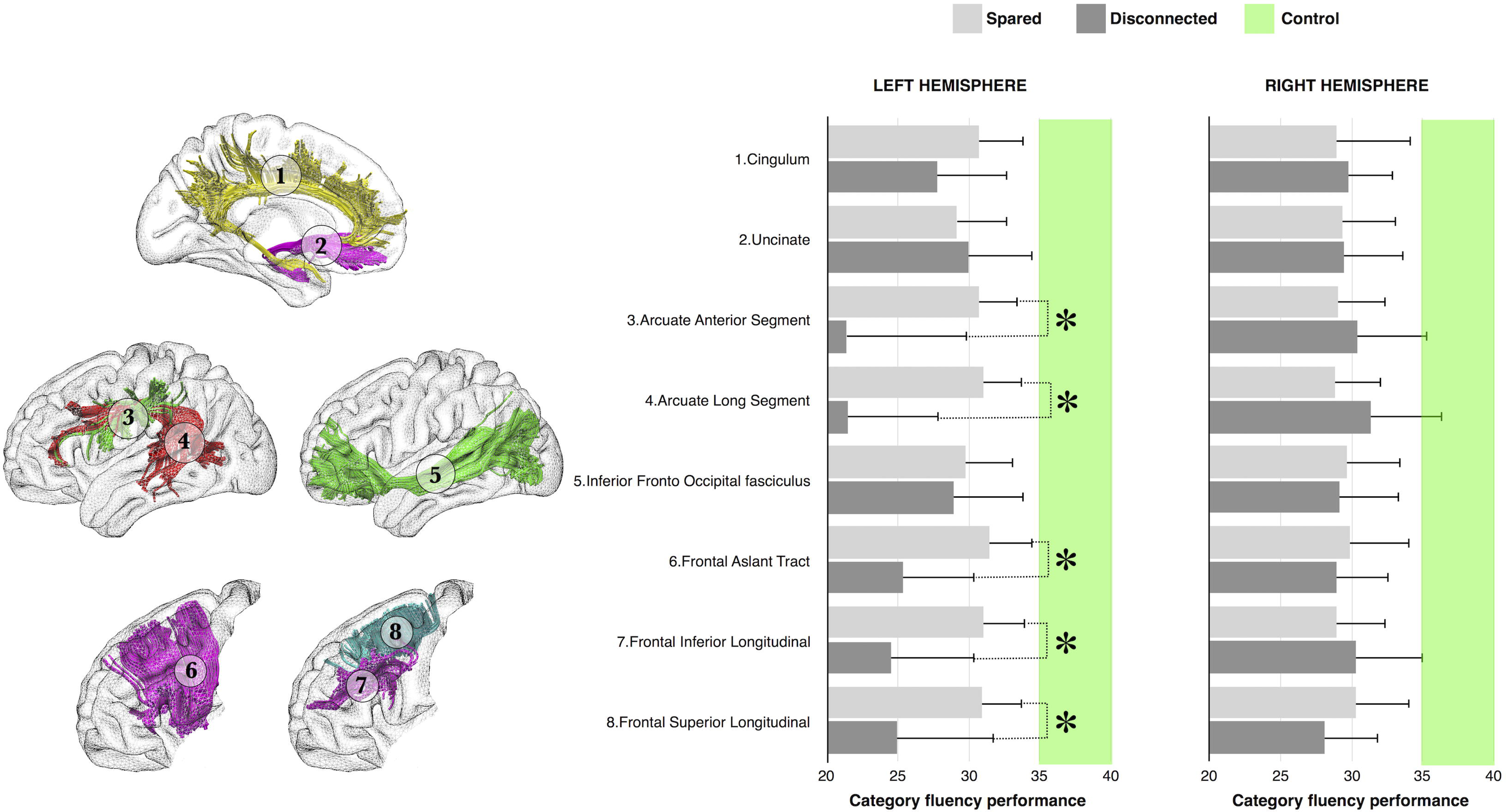
Category fluency performance (mean performance with 95% confidence intervals) for patients with (dark grey) or without (light grey) disconnection of each tract of interest. The green intervals indicate the range of controls’ performance corresponding to 95% confidence intervals. * p < 0.05

**Fig. 2:**
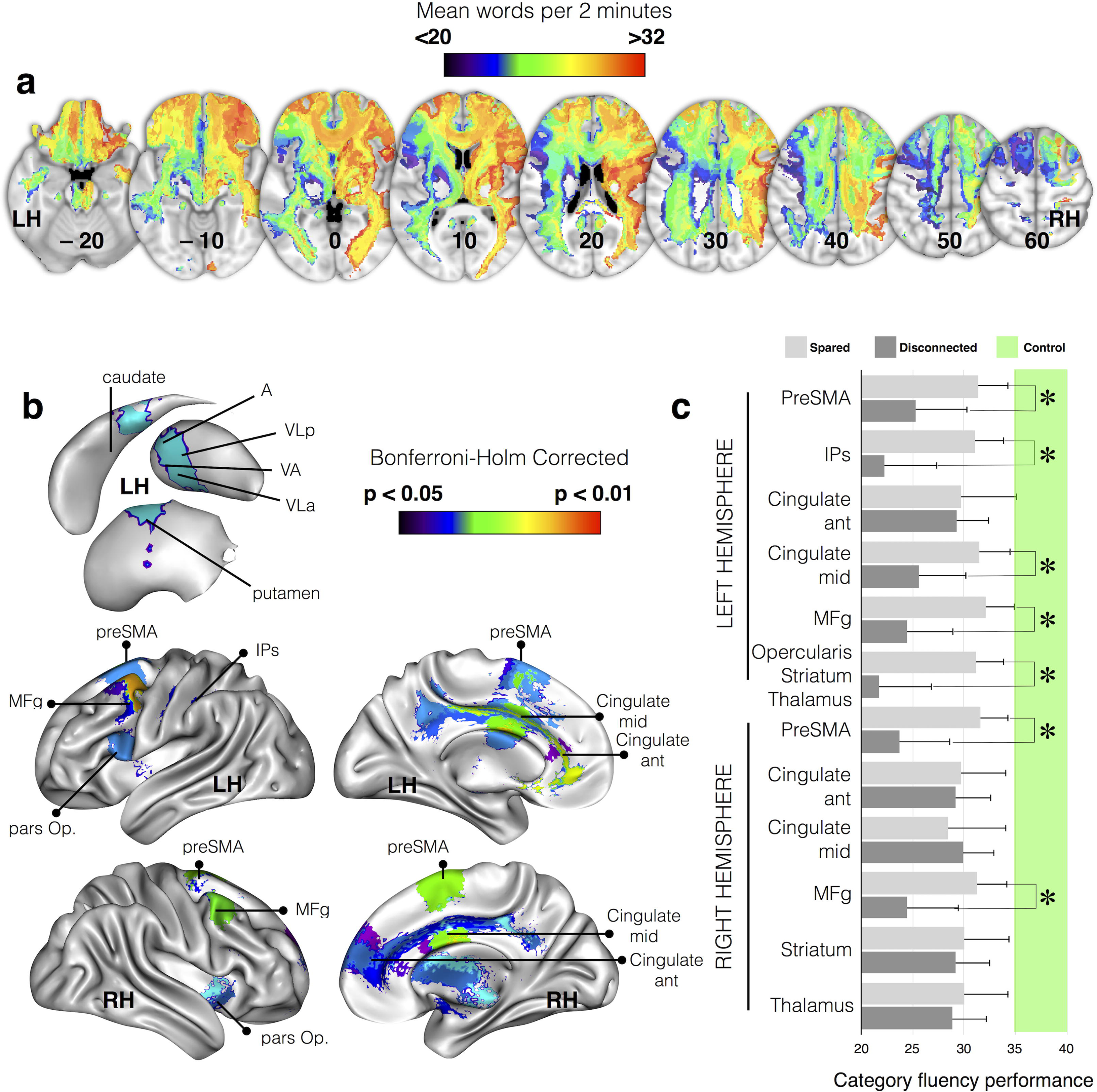
Areas directly disconnected by the lesion that significantly contributed to a decreased score on category fluency task (referred to as “disconnected areas” in the manuscript). a) Representative slices from *disconnectome maps* computed for category fluency performance, blue clusters indicate group average low performance and red high performance. b) Brain areas contributing significantly after correction for multiple comparisons. c) Category fluency performance (mean performance with 95% confidence intervals) for patients with (dark grey) or without (light grey) disconnection of each of the examined cortical regions. The green interval indicates performance in matched controls with 95% confidence intervals. preSMA: presupplementary motor area, IPs: intraparietal sulcus, MFg: middle frontal gyrus, pars Op.: frontal pars opercularis, A: anterior group of thalamic nuclei, VA ventral anterior VLp: ventrolateral posterior, VLa: ventrolateral anterior. * p < 0.05 Bonferroni-Holm corrected for multiple comparisons.

### Direct disconnection of brain areas: structural connectivity network

This approach employed the diffusion weighted imaging datasets of 10 participants in the normative group to track fibres passing through each lesion.

For each participant, tractography was estimated as indicated in [64].

Patients’ lesions in the MNI152 space were registered to each control native space using affine and diffeomorphic deformations [57], and subsequently, used as seed for the tractography in Trackvis [65]. Tractography from the lesions were transformed in visitation maps [66, 67], binarized and brought to the MNI152 using the inverse of precedent deformations. Finally, we produced a percentage overlap map by summing at each point in the MNI space the normalized visitation map of each healthy subject. Hence, in the resulting *disconnectome map*, the value in each voxel took into account the inter-individual variability of tract reconstructions in controls, and indicated a probability of disconnection from 50 to 100% for a given lesion (i.e. thus explaining more than 50% of the variance in disconnection and corresponding to a large effect size). This procedure was repeated for all lesions, allowing the construction of a *disconnectome map* for each patient/lesion. These steps were automatized in the tool *disconnectome map* as part of the *BCBtoolkit*. Note that sample size and age effects were carefully explored and reported in the supplementary material. Overall, 10 subjects are sufficient to produce a good enough *disconnectome map* that matches the overall population (more than 70% of shared variance). We also demonstrate in the supplementary material that *disconnectome maps* show a very high anatomical similarity between decades and no decrease of this similarity with age.

Thereafter, we used *AnaCOM2* available within the *BCBtoolkit* in order to identify the disconnections that are associated with a given deficit, i.e. connections that are critical for a given function. *AnaCOM2* is comparable to *AnaCOM* [68] but has been reprogrammed and optimised to work on any Linux or Macintosh operating systems.

Initially, *AnaCOM* is a cluster-based lesion symptom mapping approach, which identifies clusters of brain lesions that are associated with a given deficit, i.e. the regions that are critical for a given function. In the context of this paper, *AnaCOM2* used *disconnectome maps* instead of lesion masks, to identify clusters of disconnection that are associated with category fluency deficits, i.e. the connections that are critical for a given function. Compared to standard VLSM [8], *AnaCOM2* regroups voxels with the same distribution of neuropsychological scores into clusters of voxels. Then, for each cluster above 8mm^3^, *AnaCOM2* will perform a Kruskal-Wallis test between patients with a disconnection, patients spared of disconnection and controls. Resulting p-values are Bonferroni-Holm corrected for multiple comparisons. Subsequently, significant clusters (p-value < 0.05) are used to perform a post-hoc Mann-Whitney comparison between two subgroups of interest (i.e. disconnected patients and healthy subjects). Post-hoc results are Bonferroni-Holm corrected for multiple comparisons (statistical tests and corrections are computed using R language: [69]).

Patients-controls comparisons have been chosen as a first step in order to avoid drastic reduction of statistical power when two or more non-overlapping areas are responsible for patients reduced performance [68]. Non-parametric statistics have been chosen, as it is fair to consider that some clusters will not show a Gaussian distribution. *AnaCOM2* resulted in a statistical map that reveals, for each cluster, the significance of a deficit in patients undertaking a given task as compared to controls.

In the following sections of the manuscript, the term <u>clusters</u> systematically refers to the result of the post-hoc Mann-Whitney comparison between disconnected patients and healthy subjects that survived Bonferroni-Holm correction for multiple comparisons.

### fMRI Meta-analyses

A method described by Yarkoni et al. [70, 71] was used to identify the functional networks involved in category fluency. We searched for brain regions that are consistently activated in studies that load highly on 2 features: “fluency” (120 studies, 4214 activations) and “category” (287 studies, 10179 activations). The results were superimposed on the 3D reconstruction of the MNI152 images (**Fig. 3**).

**Fig. 3:**
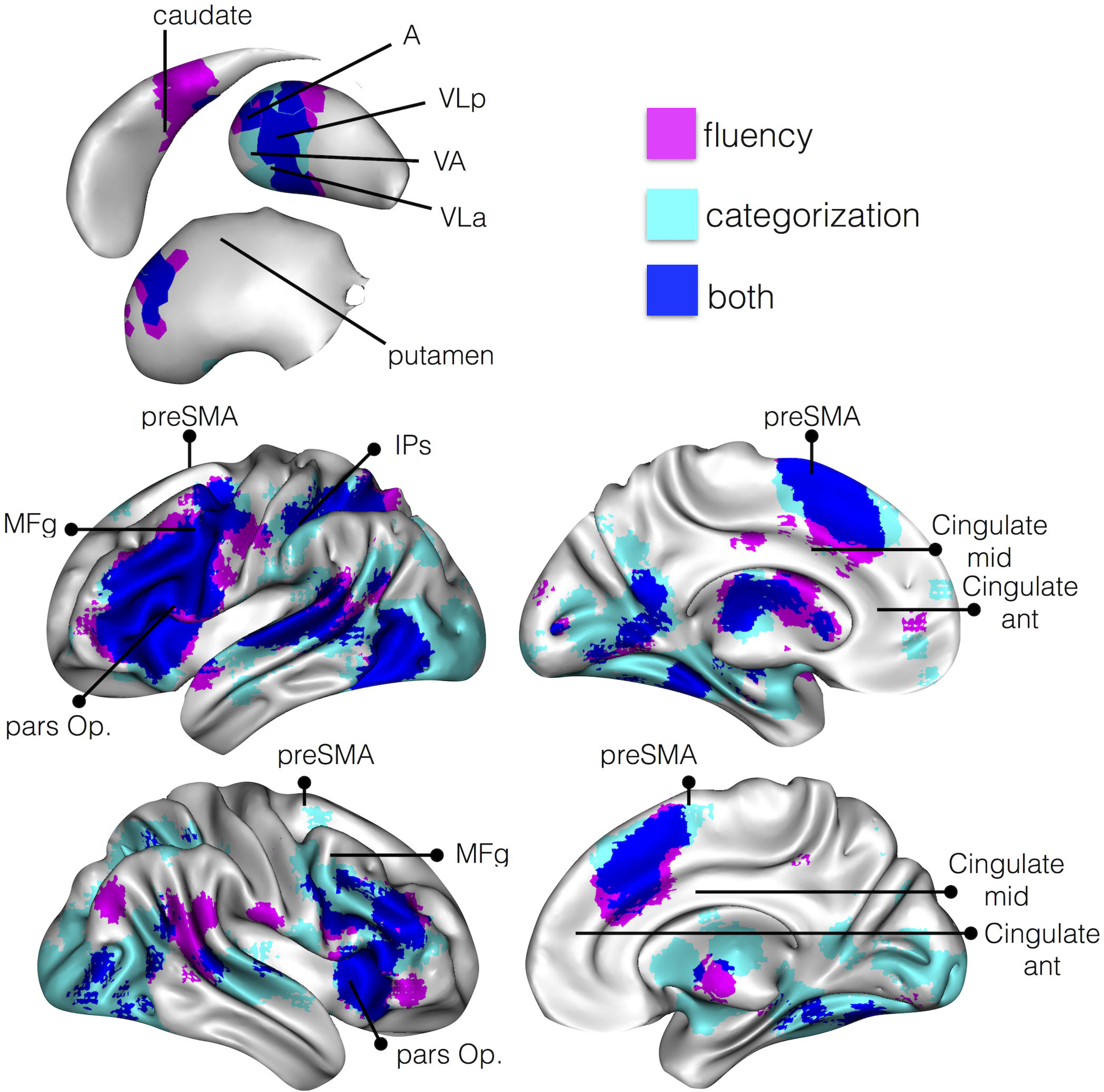
Areas classically activated with *f*MRI (p< 0.01 FDR corrected) during fluency (pink) and categorization (cyan) tasks. Areas involved in both fluency and categorization are highlighted in dark blue.

### Indirect disconnection of brain areas: functional connectivity network

Rs-*f*MRI images were first motion corrected using MCFLIRT [72], then corrected for slice timing, smoothed with a full half width maximum equal to 1.5 times the largest voxel dimension and finally filtered for low temporal frequencies using a gaussian-weighted local fit to a straight line. These steps are available in Feat as part of FSL package [73].

Rs-*f*MRI images were linearly registered to the enantiomorphic T1 images, and subsequently to the MNI152 template (2mm) using affine transformations. Confounding signals were discarded from rs-*f*MRI by regressing out a confound matrix from the functional data. The confound matrix included the estimated motion parameters obtained from the previously performed motion correction, the first eigenvariate of the white matter and cerebrospinal fluid (CSF) as well as their first derivative. Eigenvariates can easily be extracted using fslmeants combined with the –eig option. White matter and CSF eigenvariates were extracted using masks based on the T1 derived 3-classes segmentation thresholded to a probability value of 0.9, registered to the rs-*f*MRI images and binarized. Finally, the first derivative of the motion parameters, white matter and CSF signal was calculated by linear convolution between their time course and a [-1 0 1] vector.

For each control participant, we extracted the time course that corresponded to each significant cluster which was identified by the statistical analyses of the *disconnectome maps*. These time courses were subsequently correlated to the rest of the brain so as to extract seed-based resting-state networks. In order to obtain the most representative networks at the group level, for each seed-based resting-state network, we calculated the median network across the group. The median network resulting from a seed contains, in each voxel, the median of functional connectivity across all the control subjects. Medians were chosen instead of average as they are less sensitive to outliers and are more representative of the group level data [74]. The calculation of the functional connectivity was automatized and made available inside the *funcon* tool as part of *BCBtoolkit*. Medians were calculated using the function fslmaths.

Visual inspection revealed that several of these resting state networks shared a very similar distribution of activations. Therefore, an ‘activation’ matrix was derived from the seed-based resting-state networks. This matrix consisted of columns that indicated each seed-based resting-state network, and rows that represented the level of activation for each voxel in the cortex. This ‘activation’ matrix was entered into a principal component analysis in SPSS (SPSS, Chicago, IL) using a covariance matrix and varimax rotation (with a maximum of 50 iterations for convergence), in order to estimate the number of principal components to extract for each function. Components were plotted according to their eigenvalue (y) (Lower left panel in **Fig. 4**) and we applied a scree test to separate the principal from residual components. This analysis revealed that three factors were enough to explain 82% of the variance of the calculated seed-based resting-state networks. This means that three factors are good enough to summarise most of the seed-based resting-state networks results. Finally, brain regions having a statistically significant relationship with the three components (i.e. factor-networks) were detected using a linear regression with 5.000 permutations, in which the eigenvalues of the three components represented the independent variable and the seed-based resting-state networks the dependent variable. Results were Family Wise Error (FWE) corrected for multiple comparisons, and projected onto the average 3D rendering of the MNI152 template in the top panel of **Fig. 4**. In the following sections of the manuscript, the term <u>factor-networks</u> systematically refers to brain regions having a statistically significant relationship with the three components.

**Fig. 4:**
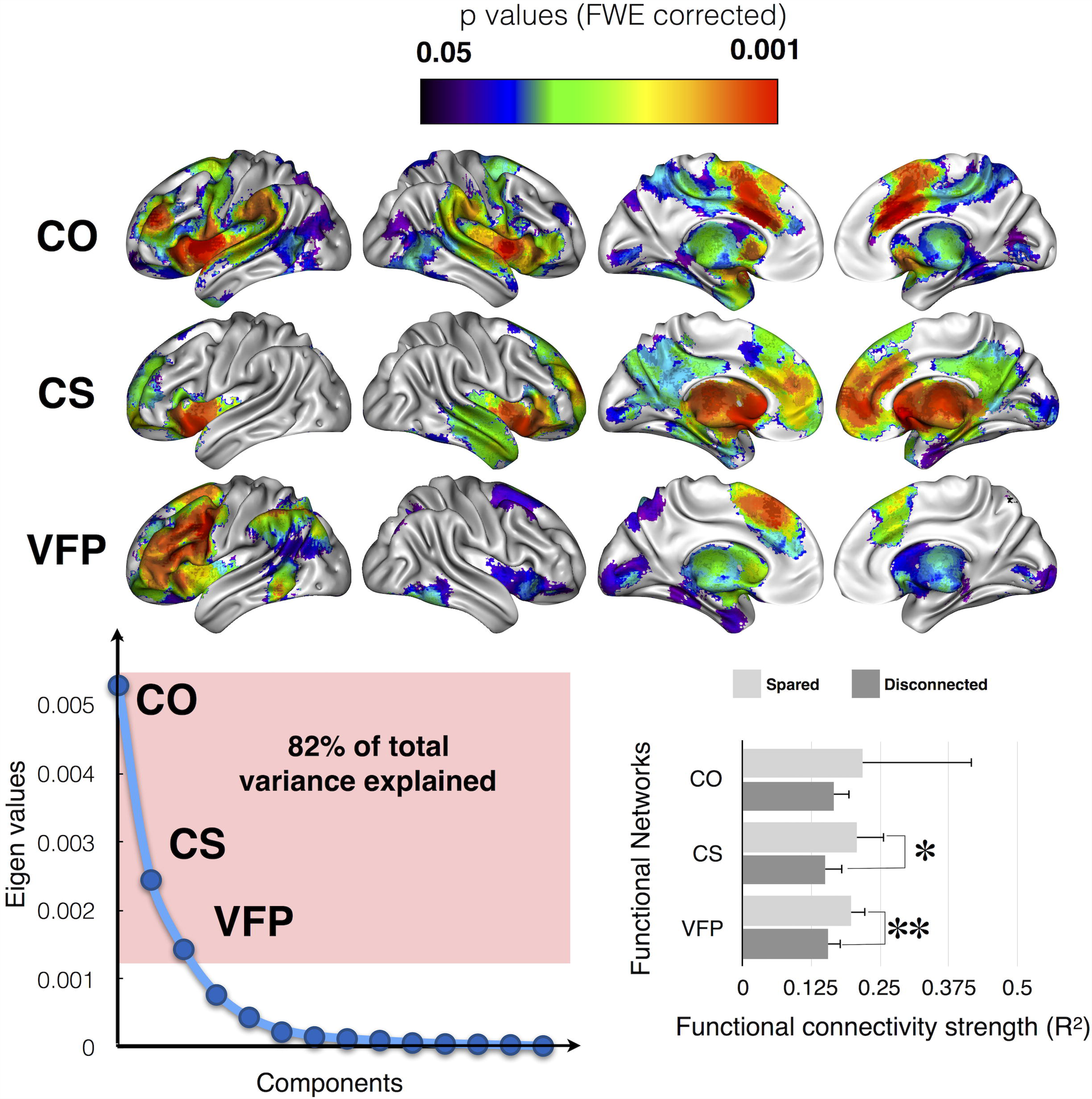
Functional networks involving the identified disconnected areas, as defined by resting state functional connectivity. Top panel, main cortical networks involving the disconnected areas revealed by a principal component analysis. Bottom left panel, principal component analysis of the raw functional connectivity result. Bottom right panel, strength of the functional connectivity for patients with (dark grey) or without (light grey) involvement of the functional network. CO: Cingulo-opercular network, CS: cortico-striatal network, VFP: Ventral fronto-parietal network. * indicates p < 0.05; **, p < 0.01

Additionally, for each patient, we extracted the time course that corresponded to each factor-network. These time courses were subsequently correlated to the rest of the brain so as to extract seed-based factor-networks in each patient. FSLstats was employed to extract the strength of factor-networks functional connectivity and subsequently, to compare patients according to their disconnection status. Note that a patient disconnected in a factor-network is a patient who has a disconnection in at least one of the cluster that contributed significantly to the factor-network.

### Structural changes in disconnected regions

A distant lesion can affect cortical macro and microstructure remotely. Conscious of this, we attempted to estimate these structural changes and their relationship with category fluency within each functional factor-network. To this aim, we explored the properties of each functional network using two complementary measures: T1w-based cortical thickness to identify fine local volumetric changes and the Shannon entropy of rs-*f*MRI as a surrogate for the local complexity of the neural networks [75]. Each original functional network seeded from each cluster was thresholded and binarized at r>0.3 and used as a mask to extract cortical thickness and entropy. Patients’ lesions were masked out for these analyses.

For the cortical thickness, a registration-based method (Diffeomorphic Registration based Cortical Thickness, DiReCT) was employed [76] from the T1-weighted imaging dataset. The first step as for the *normalisation* was to produce an enantiomorphic filling of the damaged area in order to avoid the analysis to be contaminated by the lesioned tissue. The second step of this method consisted in creating two two-voxel thick sheets, one laying just between the grey matter and the white matter and the second laying between the grey matter and the CSF. Then, the grey/white interface was expanded to the grey/CSF interface using diffeomorphic deformation estimated with ANTs. The registration produced a correspondence field that allows an estimate of the distance between the grey/white and the grey/CSF interfaces, and thus corresponded to an estimation of cortical thickness. Voxels belonging to the lesion were subsequently removed from the cortical thickness maps (see **supplementary figure 2**). This approach has good scan-rescan repeatability and good neurobiological validity as it can predict with a high statistical power the age and gender of the participants [77] as well as atrophy following brain lesions [78]. Note that the striatum and the thalamus were excluded from the cortical thickness analysis since they do not have a cortical ribbon.

Shannon entropy is an information theory derived measure that estimates signal complexity [79, 80]. In the context of rs-*f*MRI, the entropy measures the local complexity of the Blood Oxygen Level Dependent (BOLD) signal as a surrogate of the complexity of the spontaneous neuronal activity [81, 82]. Since “cells that fire together wire together” [83], for each grey matter voxel Shannon entropy of rs-fMRI can be considered as a surrogate for the complexity of the connections within this voxel and between this voxel and the rest of the brain. Shannon entropy was extracted from the previously preprocessed rs-*f*MRI using the following formula: -sum(p*log(p)) where p indicates the probability of the intensity in the voxel [75].

FSLstats was employed to extract the average cortical thickness and resting state *f*MRI entropy for each cluster and factor-network. Statistical analysis was performed using SPSS software (SPSS, Chicago, IL). In our analysis, Gaussian distribution of the data was not confirmed for the cortical thickness and the entropy measures using the Shapiro–Wilk test. Therefore, non-parametric statistics were chosen to compare cortical thickness and entropy levels between patients disconnected, spared and controls in each cluster and factor network. Additionally, bivariate Spearman rank correlation coefficient analyses were performed between the cortical thickness or entropy measurement of each functional network and each patient’s category fluency performance. Correlation significant at p < 0.0041 survives Bonferroni correction for multiple comparisons (12 networks).

## Results

### White matter tracts disconnection

Patients’ lesions were compared to an atlas of white matter connections in order to identify the probability of tract disconnections [58]. A Kruskall-Wallis test indicated that for each tract, patients (i.e. connected and disconnected) and control participants showed a significantly different performance on the category fluency test (all p < 0.001, full statistics reported in **Table 2**). Between patients, post-hoc comparisons revealed that disconnections of the left frontal aslant (U = 90.0; p = 0. 0389), frontal inferior longitudinal (U = 69.0; p = 0. 0216) and frontal superior longitudinal (U = 75.0; p = 0. 0352) tracts, the anterior (U = 28.5; p = 0. 0116) and long segment (U = 31.5; p = 0.0059) of the arcuate fasciculus were associated with a poorer performance in category fluency (**Fig. 1**). However, these post-hoc comparisons did not survive Bonferroni-Holm correction for multiple comparisons.

These results indicate that poor performance measured in patients with brain damage can be associated to some extent with white matter tract disconnections.

**Table 2:** White matter tracts disconnection relationship with category fluency statistical report. Results are not corrected for multiple comparisons. n1, number of disconnected patients; n2, number of spared patients

| Tracts | 3 groups comparison |  | Patients disconnected and connected |  | Patients disconnected and controls |  | Patients connected and controls |  | n1 | n2 |
| --- | --- | --- | --- | --- | --- | --- | --- | --- | --- | --- |
|  | K | P value | U | P value | U | P value | U | P value |  |  |
| Cingulum Left | 19 | 0.0001 | 141 | 0.2035 | 189 | 0.0003 | 277 | 0.0003 | 16 | 21 |
| Cingulum Right | 19 | 0.0001 | 161 | 0.5 | 280 | 0.0001 | 187 | 0.0019 | 23 | 14 |
| Uncinate Left | 19 | 0.0001 | 148 | 0.3994 | 176 | 0.0027 | 291 | 0.0001 | 13 | 24 |
| Uncinate Right | 19 | 0.0001 | 167 | 0.4635 | 209 | 0.0004 | 258 | 0.0003 | 17 | 20 |
| Arcuate Anterior Segment Left | 22 | 0.0000 | 29 | 0.0116 | 12 | 0.0004 | 454 | 0.0001 | 5 | 32 |
| Arcuate Anterior Segment Right | 19 | 0.0001 | 126 | 0.3855 | 118 | 0.0025 | 348 | 0.0001 | 16 | 21 |
| Arcuate Long Segment Left | 23 | 0.0000 | 32 | 0.0059 | 13 | 0.0001 | 453 | 0.0002 | 6 | 31 |
| Arcuate Long Segment Right | 19 | 0.0001 | 107 | 0.2559 | 117 | 0.0068 | 349 | 0.0001 | 9 | 28 |
| Inferior Fronto Occipital fasciculus Left | 19 | 0.0001 | 165 | 0.5 | 196 | 0.0011 | 271 | 0.0001 | 15 | 22 |
| Inferior Fronto Occipital fasciculus Right | 19 | 0.0001 | 157 | 0.3457 | 199 | 0.0002 | 268 | 0.0004 | 17 | 20 |
| Frontal Aslant Tract Left | 21 | 0.0000 | 90 | 0.0389 | 90 | 0.0001 | 377 | 0.0004 | 11 | 26 |
| Frontal Aslant tract Right | 19 | 0.0001 | 155 | 0.3131 | 194 | 0.0001 | 272 | 0.0012 | 18 | 19 |
| Frontal Inferior Longitudinal Left | 21 | 0.0000 | 69 | 0.0216 | 54 | 0.0001 | 413 | 0.0004 | 9 | 28 |
| Frontal Inferior Longitudinal Right | 19 | 0.0001 | 140 | 0.3051 | 171 | 0.0022 | 295 | 0.0001 | 13 | 34 |
| Frontal Superior Longitudinal Left | 20 | 0.0000 | 75 | 0.0352 | 73 | 0.0004 | 393 | 0.0002 | 9 | 28 |
| Frontal Superior Longitudinal Right | 19 | 0.0001 | 129 | 0.1992 | 120 | 0.0001 | 346 | 0.0005 | 13 | 34 |

### Direct disconnection of brain areas: structural connectivity network

As different white matter atlases exist for the interpretation of the white matter tract disconnection [84], and atlas-based approaches cannot assess the disconnection of the subportion of tracts nor the involvement of multiple tracts by a lesion, data driven maps of disconnection or ‘disconnectomes’ were produced. Using tractography in a group of 10 healthy controls, the registered lesions were used as a seed to reveal white matter tracts that passed through the injured area so as to produce maps of disconnections, later referred to as *disconnectome maps*. Category fluency scores were attributed to each patient’s disconnectome map (see Fig. 2a). A Kruskall-Wallis test indicated that, for several clusters, patients (i.e. connected and disconnected) and control participants showed a significantly different performance on the category fluency test (all p < 0.001, full statistics reported in **Table 3**).

**Table 3:** Direct disconnection of brain areas relationship with category fluency statistical report. Unless specified, p values are Bonferroni-Holms corrected for multiple comparisons. n1, number of disconnected patients; n2, number of connected patients

|  |  | 3 groups comparison |  | Patients disconnected and connected (uncorrected) |  | Patients disconnected and controls |  | Patients connected and controls |  | n1 | n2 |
| --- | --- | --- | --- | --- | --- | --- | --- | --- | --- | --- | --- |
|  | Disconnected areas | Kruskal Wallis | P value | U | P value | U | P value | U | P value |  |  |
| FLUENCY SCORE LEFT HEMISPHERE | PreSMA | 21.34128456 | 0.0059 | 212 | 0.0456 | 88.5 | 0.0248 | 377.5 | 0.0329 | 12 | 25 |
|  | IPs | 23.35102548 | 0.0023 | 172 | 0.0098 | 18 | 0.0295 | 448 | 0.0324 | 7 | 30 |
|  | Cingulate ant | 18.6697471 | 0.0125 | 160.5 | 0.7452 | 304 | 0.0248 | 162 | 0.0329 | 25 | 12 |
|  | Cingulate mid | 21.52289636 | 0.0054 | 219 | 0.0464 | 95.5 | 0.0141 | 370.5 | 0.0329 | 13 | 24 |
|  | MFg | 23.12826675 | 0.0026 | 237 | 0.0103 | 81.5 | 0.0054 | 384.5 | 0.0329 | 13 | 24 |
|  | Opercularis, striatum, thalamus | 23.99647373 | 0.0017 | 179 | 0.0043 | 16 | 0.0249 | 450 | 0.0324 | 7 | 30 |
| FLUENCY SCORE RIGHT HEMISPHERE | PreSMA | 22.92724537 | 0.0028 | 208 | 0.0130 | 50.5 | 0.0138 | 415.5 | 0.0329 | 10 | 27 |
|  | Cingulate ant | 18.8698263 | 0.0125 | 185.5 | 0.6470 | 225 | 0.0330 | 241 | 0.0329 | 20 | 17 |
|  | Cingulate mid | 18.62681983 | 0.0125 | 137.5 | 0.6966 | 317 | 0.0415 | 149 | 0.0329 | 25 | 12 |
|  | MFg | 22.06856039 | 0.0042 | 196 | 0.0382 | 54 | 0.0180 | 412 | 0.0329 | 10 | 27 |
|  | Striatum | 18.604408 | 0.0125 | 154.5 | 0.8966 | 310 | 0.0313 | 156 | 0.0329 | 14 | 23 |
|  | Thalamus | 19.117101 | 0.0125 | 192 | 0.5326 | 202 | 0.0248 | 264 | 0.0329 | 14 | 23 |

Results were further statistically assessed using Mann-Whitney post-hoc comparisons in order to identify areas that, when deafferented due to a disconnection mechanism, lead to a significant decrease in performance in category fluency when compared to controls.

The following results are Bonferroni-Holm corrected for multiple comparisons. Main cortical areas in the left hemisphere included the pre-supplementary motor area (PreSMA; Cluster size = 1449; Mann Whitney U = 88.5; p = 0.025), the anterior portion of the intraparietal sulcus (Cluster size = 1143; U = 18; p = 0.030), anterior (Cluster size = 837; U = 304; p = 0.025) and the middle (Cluster size = 898; U = 95.5; p = 0.014) cingulate gyrus, the middle frontal gyrus (MFg, Cluster size = 829; U = 81.5; p = 0.005), the pars opercularis of the inferior frontal gyrus (Cluster size = 5314; U = 16; p = 0.025).

In the right hemisphere, the preSMA (Cluster size = 1050; U = 50.5; p = 0.014), the MFg (Cluster size = 552; U = 54; p = 0.018), the anterior (Cluster size = 572; U = 44.5; p = 0.009) and the middle (Cluster size = 817; U = 317; p = 0.041) cingulate gyrus were also involved (**Fig. 2b**)

Subcortical areas in the left hemisphere involved the caudate, the putamen and several ventral thalamic nuclei including the ventral anterior (VA), the ventrolateral anterior (VLa) and the ventrolateral posterior (VLp) as a part of the same cluster (Cluster size = 5314; U = 16; p = 0.025)

In the right hemisphere, the striatum (Cluster size = 527; U = 310; p = 0.031), and the ventral thalamic nuclei (Cluster size = 935; U = 202.0; p = 0.025) were also involved (**Fig. 2b**).

Additionally, between patients (i.e. connected and disconnected, uncorrected for multiple comparisons) comparisons confirmed the critical involvement of the preSMA (U = 212, p = 0.0456) the MFg (U = 237, p = 0.01), the pars opercularis (U = 179, p = 0.004) and the intra-parietal sulcus (IPs, U = 172, p = 0.01) in the left hemisphere. The preSMA (U = 208; p = 0.01) and the MFg (U = 196; p = 0.038) were also involved in the right hemisphere (**Fig. 2c**).

Full statistics are reported in **Table 3**

### fMRI Meta-analyses

We further examined whether the disconnected areas in patients with poor performance are functionally engaged in tasks related to fluency and categorization using a meta-analysis approach [70, 71].

The result indicates that disconnected areas reported as significantly contributing to category fluency performance in patients are classically activated by functional MRI tasks requiring either fluency or categorization in healthy controls (**Fig. 3**).

### Indirect disconnection of brain areas

As the *disconnectome mapping* method cannot measure the indirect disconnection produced by a lesion (i.e. it fails to measure the disconnection in a region that is not directly, anatomically connected to a damaged area, but that nonetheless remains a part of the same large network of functionally connected areas), we therefore employed functional connectivity in healthy controls. This allowed us to reveal the entire network of regions that are functionally connected to the areas that were reported as contributing significantly to the category fluency performance when directly disconnected. When compared to tractography, functional connectivity has the added advantage of revealing the areas that contribute to the network through both direct, as well as indirect, structural connections.

Principal component analysis indicated that the significant areas contributing to category fluency performance belonged to 3 main functional networks (i.e. factor networks) (**Fig. 4**), which accounted for more than 80% of the total variance of the functional connectivity results.

The left cingulate clusters (anterior and middle), the right anterior cingulate, the middle frontal gyrus, the thalamus, and the operculum all belonged to the cingulo-opercular network [CO,85] including also the right preSMA, posterior cingulate and the rostral portion of the middle frontal gyrus.

The middle of the cingulate gyrus and the striatum in the right hemisphere both belonged to a cortico-striatal network [CS,86] involving the right thalamus and striatum.

Finally, the left MFg, preSMA, IPs, the pars opercularis, the thalamus and the striatum were all involved in a larger, left ventral fronto-parietal network, which also included other areas such as the right preSMA, the frontal eye field and the temporo-parietal junction [VFP,87].

Additional analyses investigated the differences in the functional connectivity of these factor-networks relative to the disconnected status of areas involved in category fluency. Between patients (i.e. connected and disconnected) comparisons revealed significantly lower functional connectivity in the left VFP network (U = 54.0, p = 0.006) and in the CS network (U = 63.0, p = 0.027) when anatomically disconnected. The CO network, however, did not show any significant difference (U = 40.0, p = 0.213). Overall, the strength of the functional connectivity for each patient did not correlate significantly with the fluency performance.

### Structural changes in disconnected regions

Additional exploratory analyses investigated structural changes related to the disconnections. We estimated these changes using two complementary measures: T1w-based cortical thickness to identify fine local volumetric changes and the Shannon entropy of rs-*f*MRI as a surrogate for the local complexity of the neural networks [75].

When compared to controls, patients showed a reduced cortical thickness in the left pars opercularis (H = 13; p = 0.0012), the MFg (H = 8; p = 0.0143), the preSMA (H = 8; p = 0.0224), the IPs (H = 9; p = 0.0131) and the right anterior (H = 7; p = 0.0296) and middle cingulate gyrus (H = 23; p = 0.000). When compared to patients with no disconnection, solely the right middle cingulate gyrus survived the Bonferroni-Holm correction for multiple comparisons (U = 67; p = 0.004). When compared to controls, disconnected patients showed reduced entropy for all regions (all p < 0.05, except for the right middle frontal gyrus). However, when compared to patients with no disconnection, none of the comparisons survived the Bonferroni-Holm correction for multiple comparisons. Uncorrected p values are reported as an indication in **table 4, and bar chart in supplementary figure 3**.

**Table 4:** Cortical thickness and *f*MRI entropy measures in disconnected areas. Uncorrected P values. n1, number of disconnected patients; n2, number of spared patients

|  |  | 3 groups comparison |  | Patients disconnected and connected |  | Patients disconnected and controls |  | Patients connected and controls |  | n1 | n2 |
| --- | --- | --- | --- | --- | --- | --- | --- | --- | --- | --- | --- |
|  | Disconnected areas | Kruskal Wallis | P value | U | P value | U | P value | U | P value |  |  |
| CORTICAL THICKNESS LEFT HEMISPHERE | PreSMA | 8 | 0.0224 | 109 | 0.0944 | 168 | 0.0057 | 514 | 0.0565 | 12 | 25 |
|  | IPs | 9 | 0.0131 | 54 | 0.0251 | 59 | 0.0019 | 667 | 0.1134 | 7 | 30 |
|  | Cingulate ant | 5 | 0.0822 | 110 | 0.1000 | 465 | 0.0175 | 269 | 0.2061 | 25 | 12 |
|  | Cingulate mid | 5 | 0.0759 | 139 | 0.2998 | 278 | 0.1436 | 435 | 0.0137 | 13 | 24 |
|  | MFg | 8 | 0.0143 | 109 | 0.0695 | 172 | 0.0028 | 502 | 0.0710 | 13 | 24 |
|  | Opercularis | 13 | 0.0012 | 40 | 0.0061 | 44 | 0.0006 | 583 | 0.0225 | 7 | 30 |
| CORTICAL THICKNESS RIGHT HEMISPHERE | PreSMA | 4 | 0.1328 | 134 | 0.4931 | 214 | 0.1711 | 523 | 0.0254 | 10 | 27 |
|  | Cingulate ant | 7 | 0.0296 | 167 | 0.4696 | 362 | 0.0191 | 295 | 0.0169 | 20 | 17 |
|  | Cingulate mid | 23 | 0.0000 | 61 | 0.0020 | 223 | 0.1414 | 254 | 0.1415 | 25 | 12 |
|  | MFg | 6 | 0.0587 | 116 | 0.2634 | 163 | 0.0359 | 538 | 0.0359 | 10 | 27 |
| SHANNON ENTROPY LEFT HEMISPHERE | PreSMA | 24 | 0.0000 | 86 | 0.2171 | 85 | 0.0004 | 210 | 0.0000 | 12 | 25 |
|  | IPs | 27 | 0.0000 | 40 | 0.0422 | 18 | 0.0002 | 260 | 0.0000 | 7 | 30 |
|  | Cingulate ant | 44 | 0.0000 | 84 | 0.1158 | 109 | 0.0000 | 3 | 0.0000 | 25 | 12 |
|  | Cingulate mid | 36 | 0.0000 | 97 | 0.3029 | 45 | 0.0000 | 127 | 0.0000 | 13 | 24 |
|  | MFg | 16 | 0.0004 | 100 | 0.4246 | 125 | 0.0043 | 272 | 0.0003 | 13 | 24 |
|  | Opercularis, striatum, thalamus | 17 | 0.0002 | 65 | 0.3680 | 75 | 0.0181 | 287 | 0.0001 | 7 | 30 |
| SHANNON ENTROPY RIGHT HEMISPHERE | PreSMA | 8 | 0.0177 | 82 | 0.2364 | 117 | 0.0078 | 413 | 0.0243 | 10 | 27 |
|  | Cingulate ant | 55 | 0.0000 | 81 | 0.0640 | 16 | 0.0000 | 4 | 0.0000 | 20 | 17 |
|  | Cingulate mid | 22 | 0.0000 | 111 | 0.4596 | 203 | 0.0001 | 114 | 0.0003 | 25 | 12 |
|  | MFg | 22 | 0.0000 | 55 | 0.0497 | 136 | 0.0533 | 209 | 0.0000 | 10 | 27 |
|  | Striatum | 23 | 0.0000 | 110 | 0.4436 | 202 | 0.0001 | 100 | 0.0001 | 14 | 23 |
|  | Thalamus | 58 | 0.0000 | 67 | 0.0204 | 0 | 0.0000 | 6 | 0.0000 | 14 | 23 |

None of these measures correlated significantly with the fluency performance.

In order to further assess the integrity of the whole network of regions that were functionally connected to the areas reported as having significantly contributed to the category fluency performance, we also extracted the cortical thickness and entropy from the regions that were functionally connected to the disconnected areas. Correlation analyses indicated that a thinner cortex in the ventral fronto-parietal network seeded from the left MFg (Spearman Rho = .464 ± 0.341; p = .004), IPs (Rho = .475 ± 0.341; p = .003) and left oper./striatum/thalamus (Rho = .512 ± 0.341; p = .001) corresponded to a reduced performance in category fluency (**Fig. 5**). Additionally, a thinner cortical thickness in the left preSMA functional network (Rho = .376 ± 0.341; p = .024) and a higher rs-*f*MRI entropy (Rho = – .420 ± 0.370; p = .019) in the mid cingulate gyrus functional network was associated with poorer performance in category fluency. These two last results, however, did not survive Bonferroni-Holm correction for multiple comparisons.

**Fig. 5:**
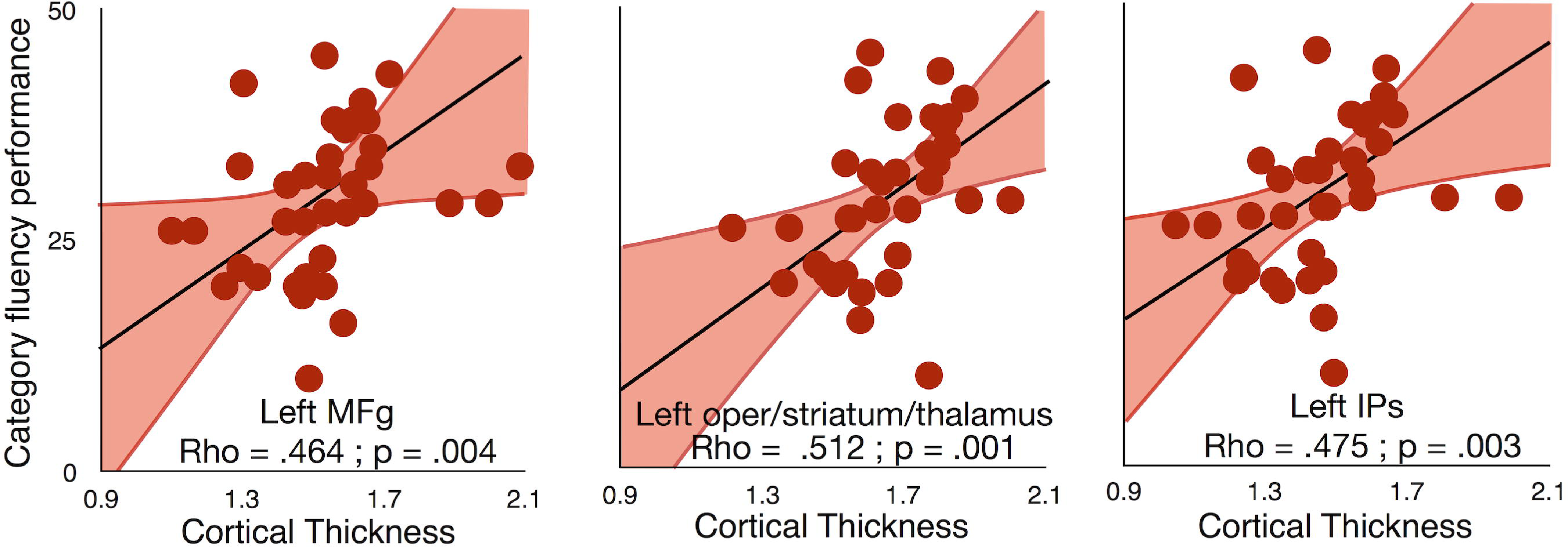
Dimensional relationship between cortical thickness measured in rs-*f*MRI disconnected networks and category fluency. Note that regression lines (in black) and intervals (mean confidence intervals in red) are for illustrative purposes since we performed a rank-order correlation.

The same analyses were repeated controlling for age and lesion size and confirmed the results for ventral fronto-parietal network seeded from the left MFg (Spearman Rho = .423; p = .01), IPs (Rho = .538; p = .001) and left opercularis (Rho = .590 ± 0.341; p < .001) corresponded to a reduced performance in category fluency (**Fig. 5**). Additionally, a thinner cortical thickness in the left preSMA functional network (Rho = .439; p = .007) and a higher rs-*f*MRI entropy (Rho = – .420 ± 0.370; p = .019) in the mid cingulate gyrus functional network was associated with poorer performance in category fluency

## Discussion

A large set of complementary methods can capture the impact of lesions on distant regions and expose the subsequent consequences upon patients’ neuropsychological performance. Several of these methods are built directly into our freely available software package BCBtoolkit. This package can be employed to measure the pathophysiological mechanisms that cause cognitive deficits, and assess the relationship between these mechanisms and their consequential effects. Here we evaluated the risk of disconnection of classically defined white matter tracts and tested their relationship with category fluency performance. We then employed a tractography-based approach in order to reveal regions that were structurally disconnected by the lesion and assess their relationship with category fluency performance as compared to controls and other patients. Functional connectivity from the disconnected regions revealed large networks of interconnected areas. Within these regions/networks, measures of cortical thickness and of entropy of the rs-*f*MRI images were correlated to fluency performance, suggesting that some structural changes that occurred within these networks were due to the remote effect of a lesion that led to cognitive impairments. Consequently, the *BCBtoolkit* provided investigators with an ability to quantify the effect of brain damage upon the whole-brain, and explore its relationship to behavioural and cognitive abilities.

The investigation into the contribution of white matter tract disconnection is more than a century old approach and postulates an interruption in the course of white matter tracts in single case patients [88, 89]. Our method provides an anatomical rationale, as well as puts forth a statistical methodology enabling it to be extended to group-level studies. In the case of category fluency performance, this analysis particularly revealed a significant involvement of the anterior and long segments of the arcuate fasciculus, which are implicated in the language network [89-91]. However, these tracts have been defined by their shape for convenience (e.g. uncinate for hook-shaped connections or arcuate for arched-shaped connections) and should not be considered as a single unit, as ultimately, sub-portions could contribute differently to the elaboration of the cognition and behaviour.

Data driven maps of disconnection or ‘disconnectomes’ were consequently produced in order to identify the sub-portion of disconnected tracts and reveal the pattern of cortico-subcortical areas that were disconnected by the lesion. For the first time, we exemplify that this method can go beyond assessing only lesions, and can be employed to assess the relationship between disconnected areas and the patient’s neuropsychological performance. Here, this approach revealed that category fluency performance significantly decreased when several cortical and subcortical clusters were directly disconnected. The observed areas are consistent with previous lesion studies on fluency tasks [92]. Furthermore, each area identified as significantly involved in this analysis corresponded, almost systematically, to activation loci derived from *f*MRI studies in healthy controls performing fluency and/or categorisation tasks. This result suggests that the method appropriately identified altered functional networks contributing to the category fluency test. Nonetheless, one might argue that a cascade of polysynaptic events can influence behaviour and that dysfunctional, disconnected areas will also impact other indirectly connected areas.

In order to explore this additional dimension, we calculated the functional connectivity of the previously identified disconnected regions (i.e. clusters). In the case of the present analysis on category fluency performance, we revealed that the disconnected areas belonged to 3 large functional networks (i.e. facto-networks): a left dominant ventral fronto-parietal network, a mirror of the right-lateralized ventral attention network [93], which link key language territories [87] and is associated with executive functions [94, 95]. We additionally showed the involvement of the cingulo-opercular network, a network that interacts with the fronto-parietal control network for the control of goal-directed behaviours [96], which together with cortico-striatal network may also be linked to a reduced performance in fluency tasks [97]. The cingulo-opercular and cortico-striatal networks may also have contributed to performance through the global inertia or the ability of participants to allocate and coordinate resources during the task [98]. Finally, disconnection was associated with a significant reduction of functional connectivity in 2 out of the 3 factor-networks investigated. This is an important result, as functional connectivity appeared to be less significantly impaired in bilateral networks, suggesting that the proportion of the preserved functional network in both of the intact hemispheres may contribute to the strength of functional connectivity.

Changes in connectivity should induce changes in the microstructure of the areas of projection, and provoke cognitive or behavioural consequences. Measures of the cortical thickness revealed a significant thinning for some, but not all, directly disconnected areas. This result may reflect a potential transneuronal degeneration mechanism [42]. However, current limitations in spatial resolution and magnetic resonance imaging signal might have biased this measure in some regions due to changes in myelination in the lower layers of the cortex [99]. Cortical thickness analyses revealed that the left dominant ventral fronto-parietal network, whether it is seeded from MFg, IPs or subcortical structures in the left hemisphere, had a reduced cortical thickness associated to the category fluency performance. This result indicates a strong and encouraging relationship between the integrity of a network derived from measures of cortical thickness and behavioural performances. Future research can benefit from this approach to stratify patients’ population and predict potential recovery. Additionally, we explored whether structural changes such as other neural (e.g. synaptic plasticity) or non-neural factors (e.g. altered properties of the vasculature) could also be captured by measures of rs-*f*MRI entropy. Our results replicated recently published results, showing a strong decrease of entropy in both hemispheres when patients were compared to controls [100]. This indicates a large-scale effect of brain lesion on the overall blood oxygen level dependent dynamic of the brain. Finally, the result between patients (connected and disconnected patients) did not survive the correction for multiple comparisons, suggesting that, although promising, Shannon entropy measures of BOLD may be too noisy of a measure to capture very fine microstructural events with high enough statistical power.

Previous reports indicated that *AnaCOM* suffers from lower specificity than VLSM (Rorden et al., 2009). *AnaCOM* compares patients with controls performances, an approach that has previously been criticised [101]. In the context of our study, classical VLSM did not reveal any significant area involved with category fluency. In classical VLSM approaches, non-overlapping lesions are competing for statistical significance, fundamentally assuming that a single region is responsible for the symptoms. In the present study, we follow Associationist principles [102, 103] assuming that several interconnected regions will contribute to the elaboration of the behaviour. By comparing the performance between patients and a control population using *AnaCOM2*, several non-overlapping regions can reach significance, without competing for it. Hence, our results differ theoretically and methodologically from previous approaches. Perhaps more importantly, the network of disconnected areas revealed by *AnaCOM2* is typically considered as functionally engaged for fluency and for categorization in healthy controls.

Newer multivariate methods have also been shown to provide superior performance compared to traditional VLSM [i.e. support vector regression lesion-symptom mapping, 7, 104]. For instance, such approaches have been employed to model the statistical relationship between damaged voxels in order to reduce false positives. In the *disconnectome maps*, this relationship has been pre-established using an anatomical prior derived from tractography in healthy controls. Therefore, it is not recommended to use multivariate approaches with the *disconnectome maps*, as they might come into conflict with the prebuilt anatomical association between the voxels. Additionally, these approaches require a much larger database of patients than the current study. Future research using large lesion databases will be required to explore the effect of multivariate statistical analysis on *disconnectome maps*.

Multivariate approaches also elegantly demonstrated that false positives can be driven by the vascular architecture [7]. This is an important limitation concerning any voxel and vascular lesion symptom mapping. Here, the group of patients explored included stroke and surgical lesions. Although we cannot exclude the participation of the vascular architecture in the present findings, the heterogeneity of the lesion included in our analyse may have limited this factor. Additionally, the statistical interaction between vascular architecture and the *disconnectome map* results remain to be explored in large database of lesions.

Methods used to estimate cortical thickness has previously been reported to perform poorly in peri-infarct regions, and the quality of the tissue segmentation may be particularly poor for stroke patients [78]. Here, we followed previously published recommendations for applying DiReCT [76] to the data from stroke patients: the lesion was masked out, the tissue segmentations were visually inspected, and manual boundary correction was performed when necessary (see **supplementary figure 2** for an example).

Finally, we applied our methods to the neural basis of category fluency as a proof of concept. The anatomy of category fluency should be, ideally, replicated in a larger sample of patients including adequate lesion coverage of the entire brain to provide a more comprehensive understanding of category fluency deficit after a brain lesion. While gathering such a large dataset of patients with brain lesions would have been impossible to achieve before, it might soon become possible thanks to collaborative initiatives such as the Enigma Consortium stroke recovery initiative (http://enigma.ini.usc.edu/ongoing/enigma-stroke-recovery/) [105].

## Conclusion

Overall, using *BCBtoolkit*, researchers and clinicians can measure distant effects of brain lesions and associate these effects with neuropsychological outcomes. However, our methods require the manual delineation of lesion masks, automatization remaining a big challenge, especially on T1-images [105]. Taken together, these neuroimaging measures help discern the natural history of events occurring in the brain after a lesion, as well as assist in the localization of functions. These methods, gathered in the *BCBtoolkit*, are freely available as **supplementary software** at [71]

## Data availability

Patients’ lesions registered to the reference map MNI152 are available as supplementary material via the following link: [106], and via the Gigascience database GigaDB [107]. However, we are not able to fully share the actual clinical sample data because sharing of the clinical raw data is not covered by the participants’ consent. A copy of the consent form as signed by the participants is available via GigaDB.

An archival copy of the supporting source code is also available via GigaDB

## Availability of supporting source code and requirements

- Project name: e.g. BCBtoolkit
- Project home page: http://toolkit.bcblab.com
- Operating system(s): Linux, MacOS
- Programming language: Java, Bash, R
- Other requirements: FSL, R, Python 2.7, Numpy
- License: BSD 3-Clause

## Authors contribution

C.F. implemented the methods inside the *BCBtoolkit*, performed the analyses and wrote the manuscript. L.C. created the pipeline for the preprocessing of the resting state and for the functional correlation and revised the manuscript. S.K. conceived and help to upgrade the statistical analyses. C.R. collected the neuroimaging data. M.U. and E.V recruited the subjects, collected and built the database of patients and matched healthy controls including the neuropsychological and neuroimaging data and revised the manuscript. E.V. also participated in the conception of the lesion study, and also provided funding for the database acquisition. R.L. provided funding for the study and revised the manuscript. M.T.d.S. wrote the manuscript, provided funding, conceived and coordinated the study, reviewed and collected neuroimaging data.

## Acknowledgments

We thank Lauren Sakuma, Roberto Toro, Jean Daunizeau, Emmanuel Mandonnet, Beatrice Garcin, Stephanie J. Forkel and the BCBlab and Brainhack for useful discussions. The authors also thank the participants of this study as well as Prof. Claude Adam, Dr. Carole Azuar, Dr Marie-Laure Bréchemier, Dr. Dorian Chauvet, Dr Frédéric Clarençon Dr. Vincent Degos, Prof. Sophie Dupont, Prof. Damien Galanaud, Dr Béatrice Garcin, Dr. Florence Laigle, Dr Marc-Antoine Labeyrie, Dr. Anne Leger, Prof. Vincent Navarro, Prof. Pascale Pradat-Diehl, and Prof. Michel Wager for their help in recruiting the patients. The research leading to these results received funding from the “Agence Nationale de la Recherche” [grants number ANR-09-RPDOC-004-01 and number ANR-13- JSV4-0001-01] and from the Fondation pour la Recherche Médicale (FRM). Additional financial support comes from the program “Investissements d’avenir” ANR-10-IAIHU-06.

